# ScrapPaper: A web scrapping method to extract journal information from PubMed and Google Scholar search result using Python

**DOI:** 10.1101/2022.03.08.483427

**Authors:** Mohd Rifqi Rafsanjani

## Abstract

This paper introduces a program called ScrapPaper, a simple Python script that use web-scraping method to extract journal information from PubMed and Google Scholar search results page. Currently the motivation behind the program development is trying to solve a problem to obtain scientific literatures information especially the title and link and save as a list for further use such as in meta-analysis and comparative study of literatures. ScrapPaper advantage that it is simple to use with no prior programming experience and get the results ready within minutes (depending on the total search result). Web-scrapping is a very powerful method to extract information from the web and ScrapPaper employ several server friendly approaches accessing both PubMed and Google Scholar site.

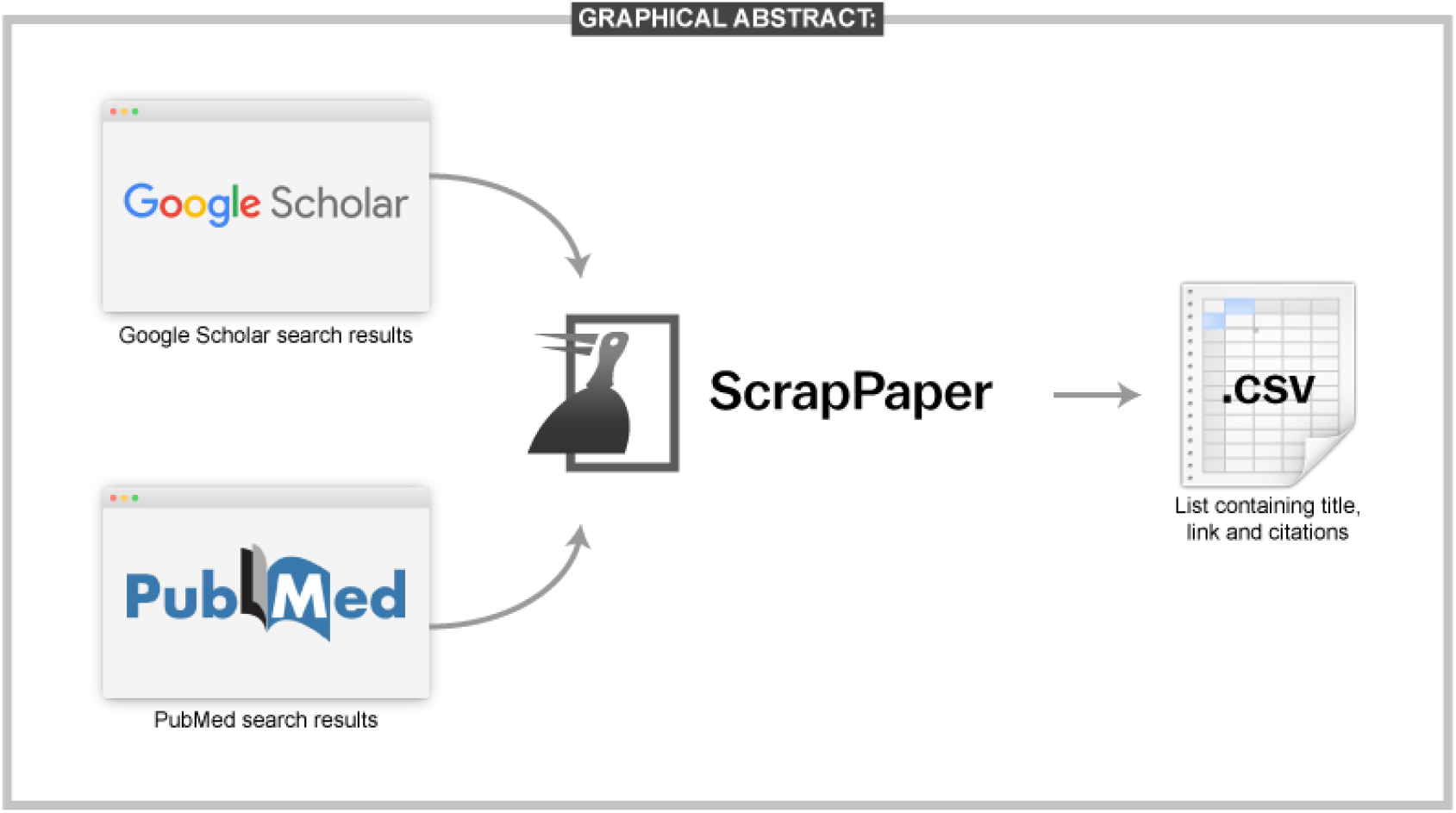

## 1 Introduction

One of the common activities carried out by researchers and scientists worldwide is finding scientific literature from databases whether via Google Scholar or any other portals such as PubMed, Semantic Scholar, Science Direct or others. Both Google Scholar and PubMed are freely publicly accessible search engine for scholarly articles and used by many especially in biomedical science field. Google Scholar, initially released in beta in 2004 by Google now contain not only peer-reviewed articles, but also patents, pre-prints, proceedings, and others [1] making its index one of the comprehensive coverages of published documents [2]. PubMed on the other hand started in 1996 developed and maintained by National Center for Biotechnology Information (NCBI) account for more than 33 million citations and abstracts focusing on biomedical literature [3].

While both platforms provide valuable tools for researchers, extracting the search results remain one of the hurdles especially for meta-analysis study or comparative study of literature. In PubMed for example, Application Programming Interface (APIs) are accessible [4] but learning to use and utilised such API required at least some background knowledge in programming language or web development. Unfortunately for Google Scholar, such API did not exist. This left researchers to manually extract the literatures information required manually either saving the page, develop some automated macros or any other copy paste method.

This paper introduces ScrapPaper, web scrapping tool to extract journal information from PubMed and Google Scholar search result developed using python. ScrapPaper basically a simple python script that only need one information of the user, which is the link of first page of the Google Scholar or PubMed search results. This simplifies many of the steps and let researchers focus on literatures rather than worrying how to obtain it. The output of the program is a .csv file that contain the list of the search results with title, link and citation of the journal or publication. Methods, guidelines and disclaimer of use are alos included in the following sections.

## 2 Methodology

### 2.1 System Requirement

- Python (version 3 or above)
- The following Python modules: *requests, csv, re, time, random, pandas, sys, bs4*
- Operating system (current code was tested on Windows 10)
- Command prompt (if using Windows)
- Search link of the first page result from PubMed or Google Scholar
- Text editor or spreadsheet software to open the results

### 2.2 Step by step instruction

#### 2.2.1 Setting up the environment

The following step applies only for first time use, once set up, simply skip these steps and continue to the following subsection (Setting up the search). If you encounter problem during this step, try running Command Prompt as administrator.

1. Install Python in your system from *https://www.python.org/downloads/*.
2. Install PIP (Pip Install Package), *https://pip.pypa.io/en/stable/installation/*. In Windows this can be achieve by simple typing the following line into the Command Prompt:
C:> py get-pip.py
3. Once PIP has been installed, using terminal (Command Prompt) download the following modules: *requests, csv, re, time, random, pandas, sys, bs4* using the following line (replace the corresponding modules name, below is the example of *requests* module):
C:> python -m pip install requests
4. Download ScrapPaper from Github via *https://github.com/rafsanlab/ScrapPaper* and extract all files into one folder.

#### 2.2.2 Setting up the search

For literature search, it is recommended to specify your search term as spesific as possible. A specific and smaller search results prevent from overloading the server and blocking you from scraping resulting faster and safer experience. Recommend to use for less than 10 pages of the search results.

1. For Google Scholar, go to *https://scholar.google.com/* and build your search terms using quotes (“…”) and/or setting up the years. Click search.
2. For PubMed, go to *https://pubmed.ncbi.nlm.nih.gov/advanced/* to build your search queries and click search.
3. Copy the link of the first page search result including *https://*.

#### 2.2.3 Running ScrapPaper

1. Open terminal (Command Prompt in Windows 10) and change directory to the location of the ScrapPaper folder using *cd* command (example):
C:> cd C:\Users\userName\Downloads\ScrapPaper
2. Run the script using the following command and wait for initialisation:
C:> python scrappaper.py
3. Once prompted, paste the first page result link copied earlier. Press Enter.
4. ScrapPaper will crawl through the search results and save the information in .csv file in the same folder. Once finish, the following line will appear:
Job finished, Godspeed you! Cite us.
5. Open the .csv file result using text editor (Notepad) or spreadsheets (Excel).

## 3 Results & Discussion

### 3.1 Example of results

Here I demonstrate one example of extracting information from Google Scholar. The search term used are: “*chromosome abnormalities*” “*chr 10*” “*chr 5*” resulting 38 search results in 3 pages. ScrapPaper is initiated following the method mentioned above and the pasted link is shown below. Extraction took around 15 seconds and the results example are shown in the following Figure 1.

**Figure 1:**
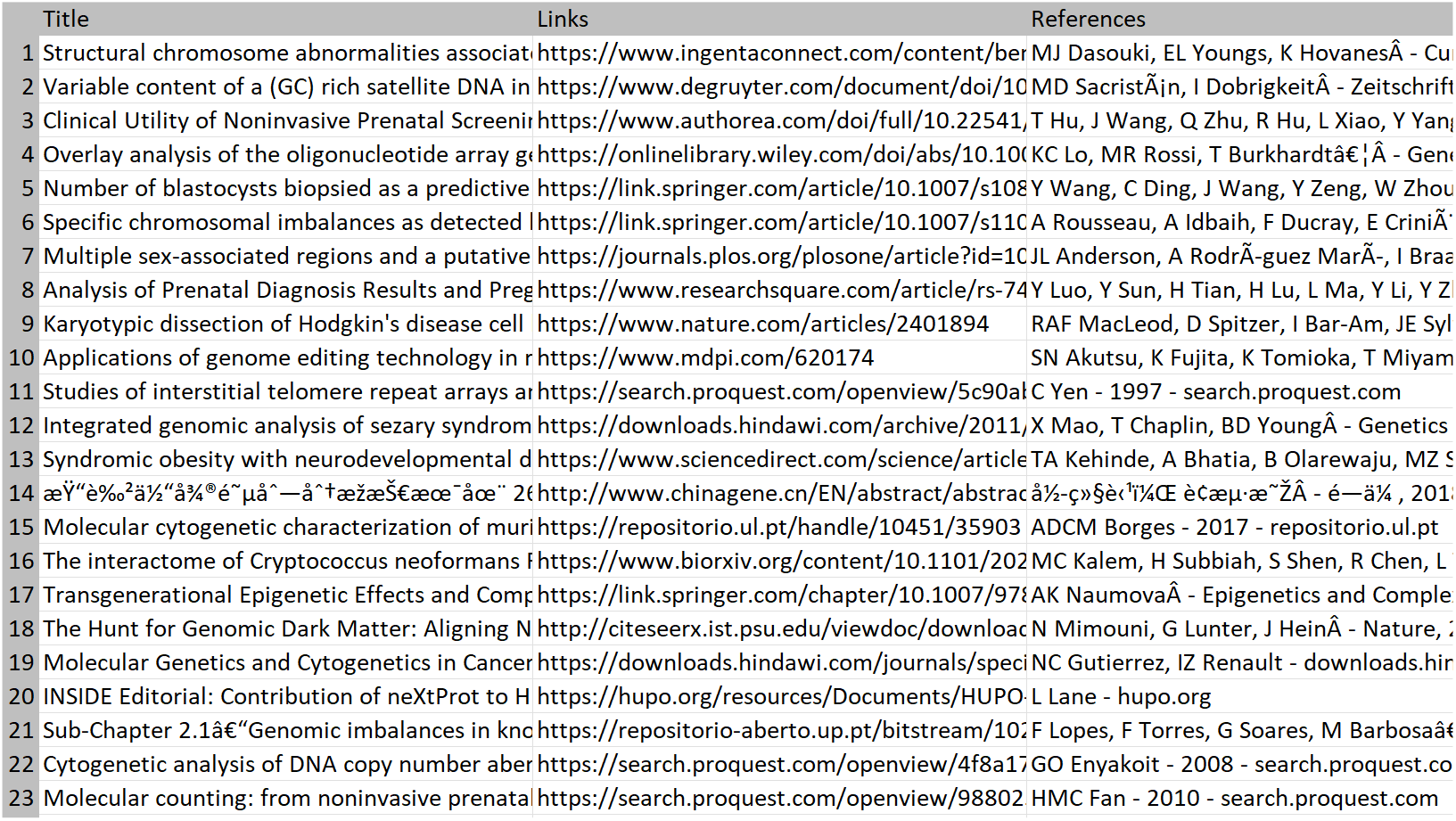
The screenshot of the .csv file of the search results opened up in Microsoft Excel.

https://scholar.google.com/scholar?l=en&as_sdt=0,5&as_vis=1&q=%22chromosome+abnormalities%22+%22chr+10%22+%22chr+5%22

### 3.2 Limitation of use

One obvious limitations of web-scrapping method is there is high chance of users getting a ReCAPTCHA to prove that it is not the bot crawling in their website. Once detected, ScrapPaper will give warning in the terminal and terminate the process immediately. This will limit the number of scraping run in one session. To avoid this, it is recommended to follow ScrapPaper guidelines mentioned in below section.

Another limitation is the incorrect display of non-English characters especially in Chinese, Russian or Japanese language due to the character encoding. Excel will display weird characters but opening in text editor like Notepad or Sublime Text will show the character just fine as they can read Unicode characters. Users are recommended to treat the .csv output file as a ‘raw file’ that can be used for further data frame manipulation in other programs.

### 3.3 Best practice & Guidelines of use

1. Specify your search as much as possible using specific search terms so that the search results only contain within a few pages rather than hundreds.
2. Avoid running multiple run in one session. This will dramatically increase your chance getting a ReCAPTCHA. Try to separate a few hours or even a day for each run.
3. Read the line in the terminal carefully, it tells you the status of the program, give you instruction what to do or even there’s an error.

## 4 Disclaimer

The method of web-scrapping using ScrapPaper is not the recommended to be use on website that do not allow robot to crawl. Hence any use of the program is on the responsibility of the user and any I as the developer of the program would not take any legal repercussion imposed by any governments, companies or any entities against the user the use the ScrapPaper and/or any modification done to it. By running the program, you already agree to above clause.

## Supporting information

Supplement Results 1

